# Identification and functional validation of human islet microRNAs associated with donor trait

**DOI:** 10.1101/2022.10.14.512222

**Authors:** Wilson K.M. Wong, Isabelle El-Azzi, Aditya Nachanekar, Ehsan Alvandi, Ho Trong Nhan Pham, Mya Sara, Feifei Cheng, Guozhi Jiang, Anja E. Sørensen, Yi Vee Chew, Thomas Loudovaris, Helen E. Thomas, Ronald C.W. Ma, Wayne J. Hawthorne, Louise T. Dalgaard, Mugdha V. Joglekar, Anandwardhan A. Hardikar

## Abstract

**Objectives:** Human islets are widely researched to understand pathophysiological mechanisms leading to diabetes. Sex, age, and body mass index (BMI) are key donor traits influencing insulin secretion. Islet function is also regulated by an intricate network of microRNAs.

**Methods:** Here, we profiled 754 microRNAs and 58,190 potential targets in up to 131 different human islet donor preparations (without diabetes) and assessed their association with donor traits. We further performed mechanistical studies to observe the causal role of the age-associated key microRNAs on relative telomere length in human islets.

**Results:** MicroRNA discovery analyses identified miR-199a-5p and miR-214-3p associated with sex, age and BMI; miR-147b with sex and age; miR-378a-5p with sex and BMI; miR-542-3p, miR-34a-3p, miR-34a-5p, miR-497-5p and miR-99a-5p with age and BMI. There were 959 mRNA transcripts associated with sex (excluding those from sex-chromosomes), 940 with age and 418 with BMI. MicroRNA-199a-5p and miR-214-3p levels inversely associate with transcripts critical in islet function, metabolic regulation, and senescence. Our functional studies verified that inhibition of these two microRNAs (miR-199a-5p/-214-3p) slowed down telomere length shortening in human islet cells maintained in vitro and demonstrating cellular senescence.

**Conclusions:** Our analyses identify human islet cell microRNAs influenced by donor traits.

**Graphical abstract:** 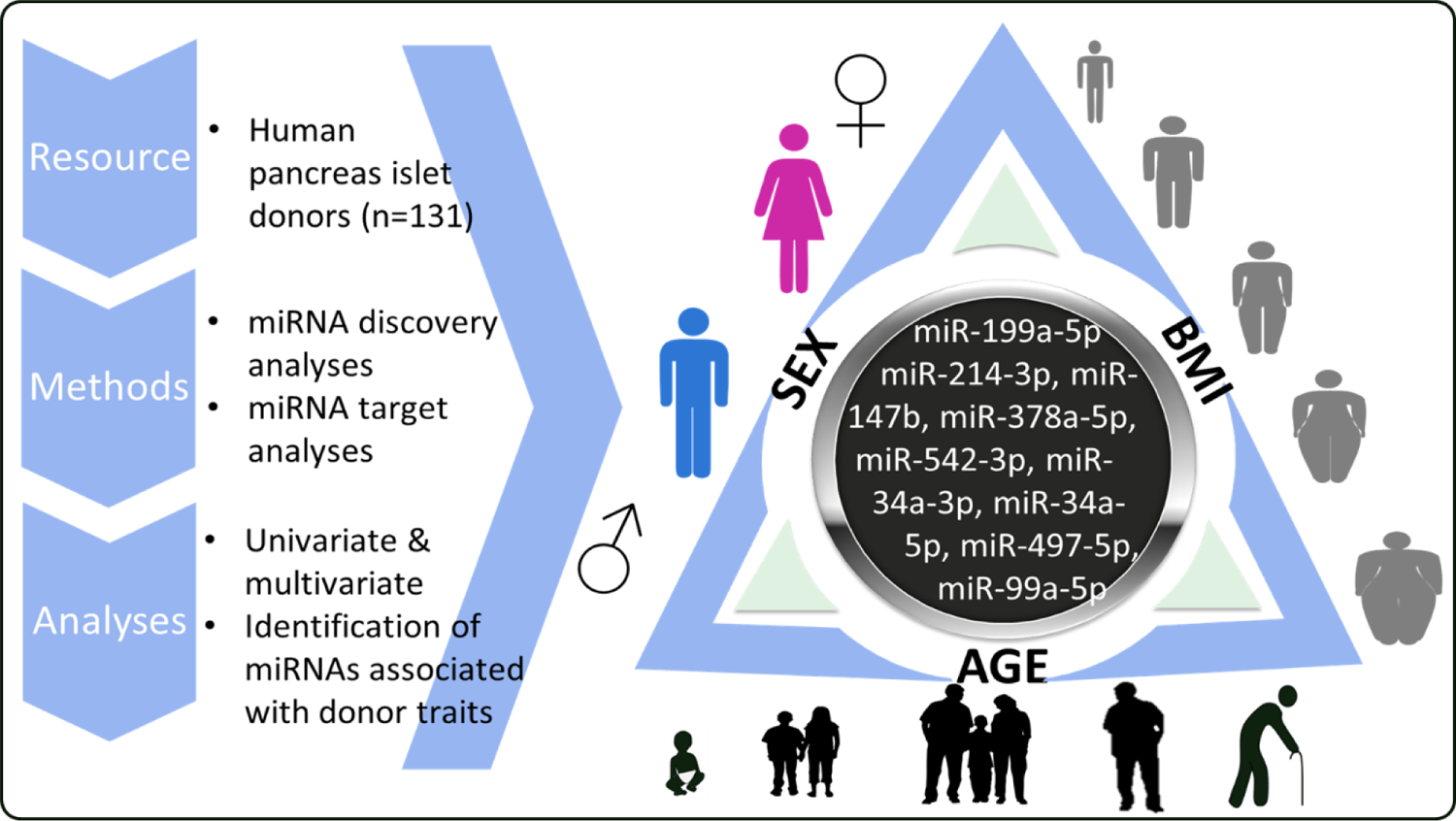

## 1. Introduction

Human pancreatic islets are widely used in research to identify/validate dysregulated metabolic or apoptotic pathways leading to beta-cell failure or death in diabetes. Sex, age, and body mass index (BMI) have been indicated to influence diabetes risk ^1–3^.

Sex-associated differences in islet glucose response, beta-cell mass, islet oxidative stress and type 1 diabetes (T1D) incidence have been reported ^4^. Testosterone, the main sex hormone in males, has been shown to enhance insulin production in mice and humans ^5,6^. DNA methylation patterns between islets of male and female donors have been linked with differences in islet gene expression, function and insulin secretion ^7^.

Age-related differences in DNA methylation are also reported in human islets ^8^. Older (>40 years old) donor islets have lower insulin secretory response to glucose compared to those from younger donors, which may be associated with changes in beta-cell ATP generation ^9^. A declining trend in beta- and non-beta-cell mass has been observed in islets from newborn to older human organ donors (aged from 0 to 79 years old) ^10^. Reductions in calcium coordination, gap junction coupling, and insulin secretion dynamics were also observed in islets from older organ donors ^11^.

Body mass index (BMI) is a measure of obesity and an important determinant of diabetes progression ^12,13^, highlighting the potential link to beta-cell function. Elevated levels of inflammatory macrophages in islets from obese individuals (BMI > 30) are contributors to insulin resistance and type 2 diabetes (T2D) susceptibility ^14^.

Previous studies have explored the differences in transcripts associated with donor sex, age, and BMI in human islets ^7,8,15–17^. However, knowledge of transcriptomic variations in human islets at non-coding RNA (such as microRNA) level due to differences in sex, age and BMI of donors is lacking. This is the first report analysing microRNAs and their targets in a large set (n=131) of human islet preparations stratified for donor traits (sex, age and BMI). Our analyses also determine correlations of microRNAs with mRNA transcripts profiled in a sub-set (n=33) of human islets. We further investigated the causal role of the two age-, sex- and BMI-associated microRNAs in ageing, as represented by replicative senescence^18^ (via multiple passaging). Inhibition of these two microRNAs slowed down telomere length shortening in human islet-derived cells. These findings emphasize the influence of donor traits on human islet microRNA expression.

## 3. Results

### 3.1. Human islet microRNAs associated with sex, age and BMI

We assessed human islet samples (n=101, **Figure 1A** for microRNAs and n=63 for mRNAs; **Figure 1B**) for their associations with donor sex, age, and BMI. Correlation analyses between selected microRNAs and mRNA datasets were performed in a subset (n=33) of islet donors (**Figure 1C**). A study schematic for analyses from both Australian islet isolation sites (at SVI and WIMR) is presented (**Figure 1D**).

**Figure 1.**
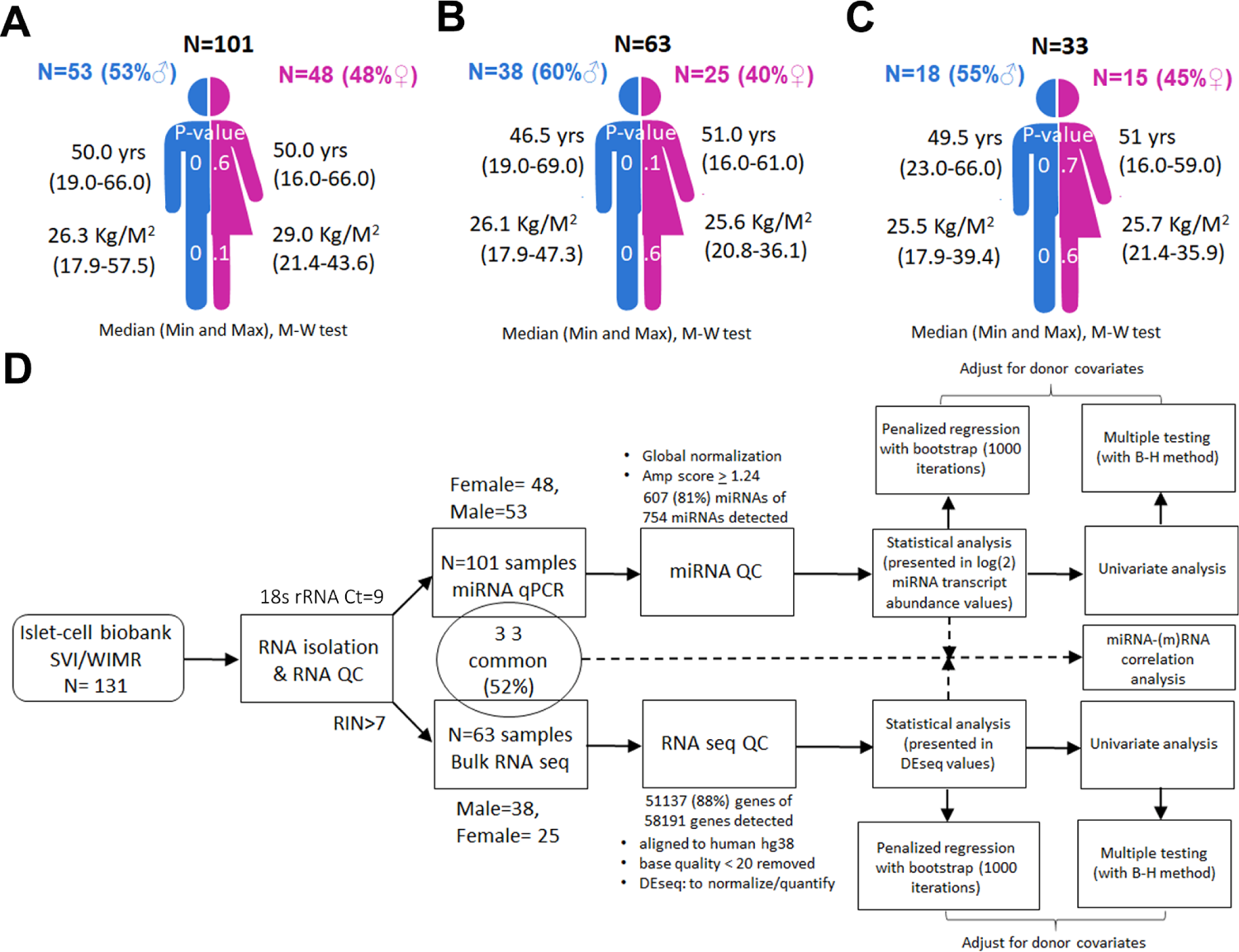
Study workflow. Summary of the cadaveric donor islet samples (**Table S1**) and their traits (sex, age and BMI) used in **(A)** microRNA analysis (n=101) and **(B)** for RNA-seq (n=63); **(C)** microRNA-mRNA correlative analysis (n=33). Mann-Whitney test was used to assess significant differences between male and female islets. The median, minimum and maximum range are presented for donor age (year) and BMI (kg/m^2^). (**D**) Flow chart of the study design. A total of 131 human islet samples obtained from different donors without diabetes through the Australian Islet Transplantation Program (at Westmead Institute for Medical Research (WIMR), Westmead Hospital, Sydney, and the St Vincent’s Institute (SVI), Melbourne) was used in this study. Samples were isolated for RNA and randomly selected for microRNA (qPCR) and/or mRNA (bulk RNA-seq) analyse. Statistical analysis of donor traits (sex, age or BMI) was performed using univariate, multiple testing (using Benjamini-Hochberg approach), adjustment to other respective traits (covariate) and machine-learning penalized regression with bootstrap analysis. Correlative analysis was performed to identify correlation between microRNA and mRNA.

We performed univariate analyses and adjustments for the other two traits (sex and/or age and/or BMI) on profiles of 754 discovery microRNAs in our set of human islet samples (n=101; **Figure 1A** and **Table S1**) to identify microRNAs with significant association with donor sex, age, and BMI (**Figure 2A and Table S2**). Machine-learning penalized regression with bootstrapping offered validation. We identified 10 microRNAs associated with sex, 32 microRNAs associated with age and 47 microRNAs associated with BMI, based on a significance cut-off (p≤0.05) after adjustment for the other remaining traits (**Figure 2A and Table S2**). Two microRNAs (miR-199a-5p and miR-214-3p) were associated with the three donor traits (sex, age and BMI), while several others (**Figure 2A and Table S2**) were associated with at least two of the donor traits and the importance of these microRNAs was verified using penalized logistic/linear regression (PLR) and bootstrapping.

**Figure 2.**
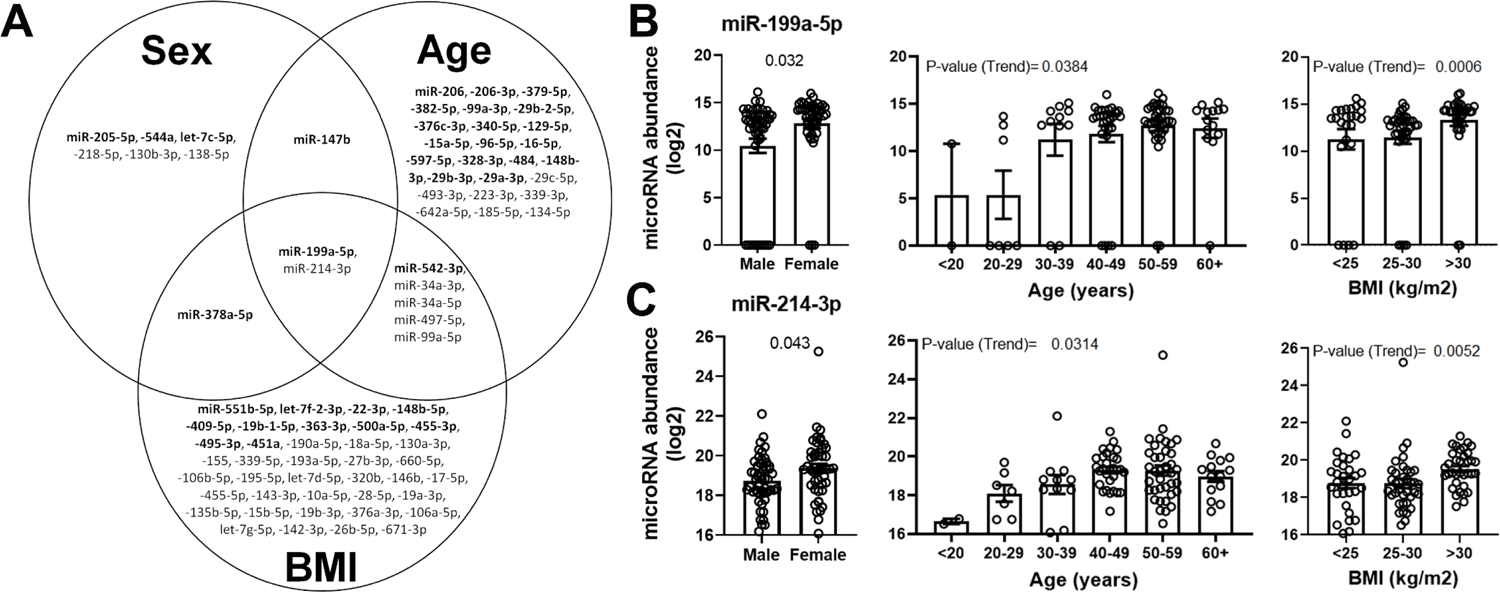
Human islet microRNAs associated with sex, age and BMI. **(A)** Venn diagram of significantly associated microRNAs with donor sex, age and/or BMI. Only the microRNA which are detectable at least 50% of the samples and have statistical significance after adjustment (to their other two respective traits) are shown. MicroRNAs which are present also in the bootstrap (revalidation on penalized regression) analyses are in bold. MicroRNAs are ordered by highest to lowest frequency in bootstrap. Real-time TaqMan® qPCR data presented in microRNA transcript abundance (log2) for **(B)** miR-199a-5p and (**C**) miR-214-3p for sex, age and BMI. The adjusted (for age and BMI) p-value between male and female (sex) comparison are presented. Trend analysis (using Kruskal-Wallis test) was performed to calculate p-value (trend) presented (in figures) for age or BMI. Mean ± SEM are presented.

Both miR-199a-5p and miR-214-3p are expressed at significantly higher levels in female compared with male islets (adjusted p-value=0.032 and 0.043 respectively) and are significantly and positively correlated with BMI (adjusted p-value=0.017 and 0.039 respectively; **Table S2**). A significant increasing trend for miR-199a-5p and miR-214-3p levels was observed for islets from lean (BMI 17.9 – 24.9), overweight (BMI 25-30) and obese (BMI >30) groups (**Figure 2B** and **2C**; trend p-values 0.0006 and 0.0052, respectively). Both microRNAs (miR-199a-5p and miR-214-3p) are significantly and positively correlated with age after adjustment for other traits (adjusted p-value=0.017 and 0.050 respectively, **Table S2**). When grouped by decades of age, a significant trend was observed for miR-199a-5p and miR-214-3p (**Figures 2B** and **C**; trend p-values 0.0384 and 0.0314, respectively).

MicroRNA-147b positively correlated (**Table S2**) with age and was detected at significantly high levels in islets from females than males (adjusted p-value=0.013, **Figure S1A, Table S2**). MicroRNA-378a-5p was significantly different between sex (adjusted p-value=0.029) and was also significantly (**Figure S1B**) and positively correlated with BMI (**Table S2**).

We found miR-542-3p, miR-34a-3p, miR-34a-5p, miR-497-5p and miR-99a-5p positively correlated (**Table S2**) with age and BMI. We observed that miR-34a-5p, miR-34a-3p and miR-497-5p show a significant increasing trend with age as well as with BMI (**Figure S1D-F**), while miR-99a-5p only showed a significant increasing trend for BMI (**Figure S1G**).

### 3.2. Target mRNAs associated with sex, age and BMI in human islets

Next, we analysed RNA-seq profiles from human islets (n=63, **Figure 1B** and **Table S1**) for associations with donor sex, age, and BMI. We identified 959 (excluding X- and Y-linked genes, **Table S3**), 940 (**Table S4**) and 418 (**Table S5**) gene transcripts associated with sex, age, and BMI respectively (p-value<0.05), after adjustment for the other two traits examined and following FDR testing. Many of these transcripts have been reported in previous publications and associated with age, sex, and BMI; while our analyses also identified new associations between some of the transcripts and donor traits (**Table S3-S5**).

### 3.3. Expression quantitative Trait Loci (eQTLs) present in sex-, age- and BMI-associated genes in human islets

Candidate genes significantly associated with sex, age or BMI (**Tables S3-S5**) were assessed on the Translational human pancreatic islet genotype tissue-expression resource (TIGER, http://tiger.bsc.es/; containing RNA-sequence and genotype profiles of 404 human islets) to identify eQTLs variations in human islets ^19^. Genes such as *BARHL1*, *ADTRP* and *RASAL1* identified in our analyses have significant eQTLs variants among the islets in the TIGER database (**Table S6**).

### 3.4. Sex-, age- and BMI-associated microRNAs inversely correlate with mRNA transcripts critical in islet function, metabolic regulation, and senescence

To further explore networks between sex-, age- and BMI-associated microRNAs and mRNA transcripts, correlative analysis was performed on their expression profiles in a subset of human islet samples (n=33; n=15 female and n=18 male, **Figure 1C** and **Table S1**). Functional interactions between microRNAs and mRNAs were assessed *in silico* (TargetScan Human ver.8). We observed significant negative and positive correlations between miR-199a-5p and miR-214-3p to target mRNAs (**Table S7 and S8**). Since majority of targeting microRNAs reduce mRNA expression ^20,21^, we focussed on negatively correlated mRNAs with potential binding sites for these microRNAs. The top-10 significantly and negatively correlated mRNAs with highly conserved binding sites to miR-199a-5p are presented in **Figure 3A**, while **Figure 3B** presents the top-10 most significant negatively correlated mRNAs that contain known (albeit poorly conserved) binding sites to miR-214-3p.

**Figure 3.**
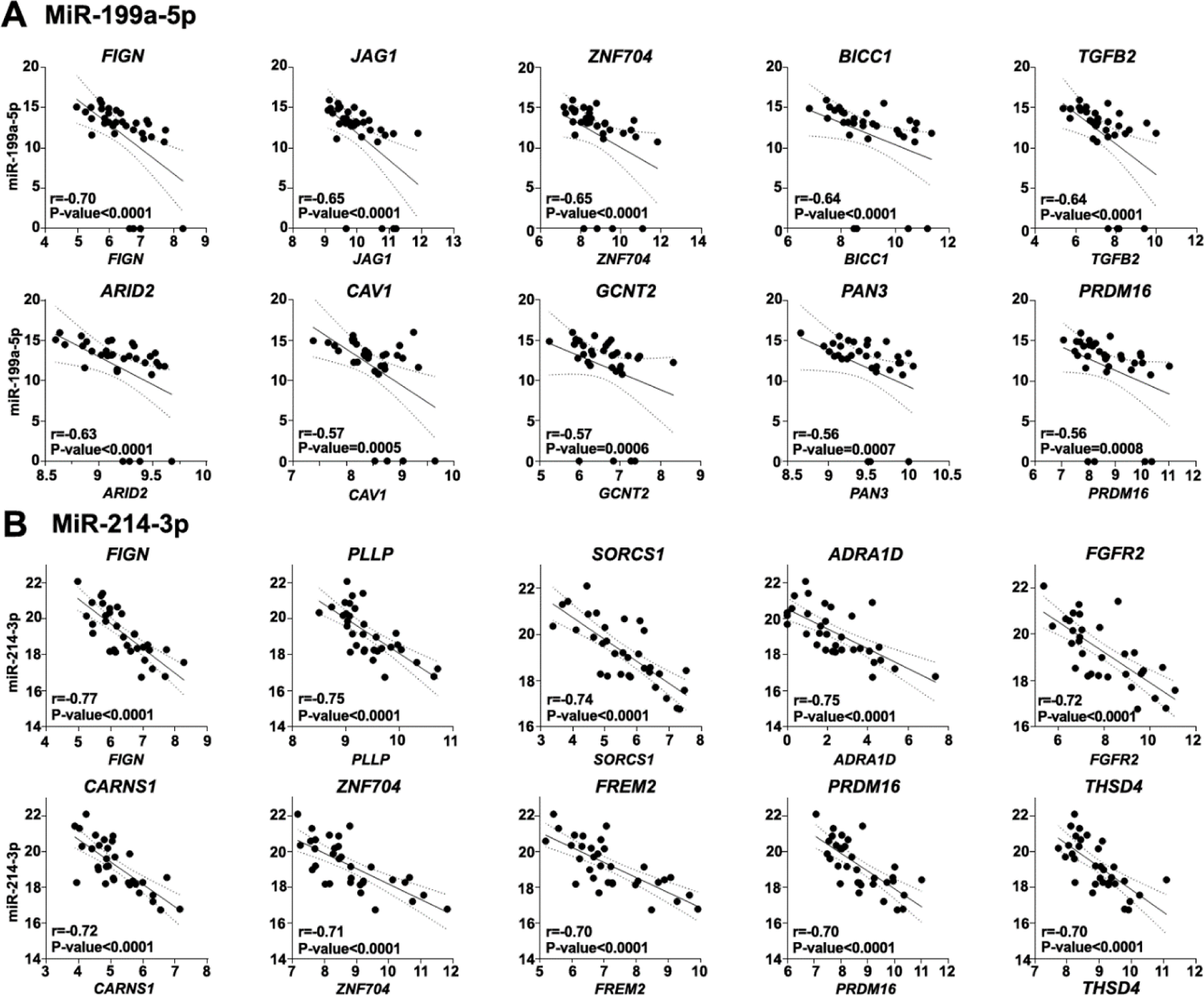
Sex, age and BMI microRNAs inversely correlate with critical islet function, metabolic regulation, and senescence mRNA transcripts in human islets. (**A**) MiR-199a-5p and (**B**) miR-214-3p correlation to their 10 most significantly negatively correlated genes, which contain potential conserved binding site (related to **Table S7 and Table S8 respectively**). Presented here are the 10 most significantly negatively correlated genes with miR-214-3p. Note that miR-214-3p target genes were had poor evolutionary conservation. The 95% confidence interval bands (dotted lines) and best-fit line (bold line) are presented. The Spearman coefficient (r, rho) and p-value are presented for each correlation plot.

MiRNA-199a-5p significantly and positively correlated to 3002 mRNAs and negatively correlated to 3136 mRNAs (**Table S7**). There were 427 genes, which negatively correlated to miR-199a-5p with potential binding sites (for miR-199a-5p), and 74 mRNAs from those had conserved binding sites to miR-199a-5p.

MiR-214-3p, significantly and positively correlated to 3358 mRNAs and negatively to 3869 mRNAs (**Table S8**). There were 815 of the 3869 negatively correlative mRNAs with binding sites for miR-214-3p.

Most of the microRNAs associated with at least two traits (miR-147b, miR-199a-5p, −214-3p, miR-542-3p, 34a-3p, −34a-5p, miR-497-5p, miR-99a-5p and miR-378a-5p; **Figure 2A**) correlated significantly with each other (**Figure S2**). We also observed common mRNA targets for these microRNAs such as *PRDM16* and *FIGN,* that significantly and negatively correlated with both miR-199a-5p and miR-214-3p (**Figure 3A** and **3B**) respectively.

### 3.5. Functional T2D risk genetic variants of mRNA transcripts associated with miR-199a-5p and miR-214-3p

The top 10 mRNAs that most significantly and negatively correlated with miR-199a-5p or miR-214-3p and contained potential binding sites for these two microRNAs were assessed to identify genetic variants linked to T2D risk traits (https://t2d.hugeamp.org/). Interestingly, *FIGN* and *PRDM16 (*associated with miR-199a-5p and miR-214-3p), *PAN3* (associated with miR-199a-5p) and *THSD4* (associated with miR-214-3p) contain genetic variants, which significantly associated with BMI (**Table S9**). *TGFB2* (associated with miR-199a-5p) and *CARNS1* (associated with miR-214-3p) also contained genetic variants significantly associated with trunk fat ratio.

### 3.6. Sex-, age- and BMI-associated islet microRNAs are involved with transcriptional, translational, cell cycling and metabolic pathways

Pathway over-representation analyses were performed on the predicted mRNA targets of microRNAs significantly associated with sex, age and BMI. Messenger RNA targets of the seven microRNAs (miR-199a-5p, −214-3p, −544a, let-7c-5p, −218-5p, −130b-3p, −147-3p and −138-5p) that have higher abundance in female islets, were associated with terms related to cell cycle and metabolic processes. Moreover, transcriptional and translational pathways such as mRNA processing, splicing, and translation initiation, as well as sex differentiation were also enriched (**Figure S3**). The two microRNAs (miR-378a-5p and −205-5p) with higher abundance in male islets, were over-represented in categories relating to generation of precursor metabolites/energy and mRNA translation. Pathway analysis on mRNAs targeted by microRNAs associated with donor BMI showed involvement in metabolic processes and biological regulatory pathways. Many of the metabolic pathways (related to BMI-associated microRNAs), and cell signalling pathways were observed to be involved with donor age-associated microRNAs. Pathway analysis on miR-199a-5p and −214-3p targets revealed that their combined mRNA targets are over-represented in cell-cell signalling, biological regulation, metabolic, mRNA and translation pathways (**Figure S3**).

### 3.7 Effect of miR-199a-5p and −214-3p knock-down on human islet telomere length

We then investigated the causal relationship between the two microRNAs (miR-199a-5p and miR-214-3p) with ageing. Expression levels of miR-199a-5p, miR-214-3p and relative telomere length (RTL) ^18^ were measured in human islet derived cells undergoing multiple (up to 16) passages *in vitro*, representing replicative cellular senescence (ageing). Statistical significance was observed between miR-199a-5p and RTL (r= −0.29, P-value= 0.0467) (**Figure S4A**). MicroRNA-199a-5p remained statistically significant to RTL after adjustment to age, sex and BMI (P-value= 0.0246). While miR-214-3p although not significant, also was negatively correlated to RTL. MicroRNA-199a-5p and miR-214-3p significantly positively correlated (P-value <0.0001) with each other (**Figure S4A**). To understand the function of these two microRNAs on RTL in replicative cellular senescence (indirect measure and representation of ageing); we performed their knock-down (with LNA power inhibitors) in human islet-derived cells that underwent multiple passaging *in vitro* (**Figure S4B**). LNA power inhibitor induced 91-99% inhibition for both microRNAs compared to the baseline expression levels. Interestingly, when both microRNAs were inhibited together during propagation of islet-derived cells *in vitro*, a 22.97 % reduction in telomere attrition was observed (**Figure S4B**).

## 4. Discussion

With increasing shortage and costs associated with human islet isolation, most studies involving human islets assess a smaller donor-set. Our analyses demonstrate that human islet microRNA (and their target-gene) expression may be significantly different due to donor characteristics and that these differentially expressed microRNAs are associated with cell cycle/signalling, mRNA translational and metabolic pathways(**Figure S3**).

Several microRNAs (such as miR-214-3p, miR-199a-5p, miR-147b, miR-542-3p, miR-99a-5p and miR-378a-5p) identified in our study, have not been previously reported to be associated with sex, age or BMI, across other human tissues, while some microRNAs (miR-34a-5p and −3p) positively correlated with age and BMI in human vascular endothelial cells^22^ and in rat beta-cells ^23^.

In our study, two microRNAs (miR-199-5p and miR-214-3p) were associated with all three donor traits. Circulating miR-199a-5p was reported to correlate with BMI and fat mass in a childhood obesity study ^24^. In addition, exosomal miR-199a-5p has been identified to be associated with hepatic lipid accumulation and fatty acid metabolism ^25^. In our analysis, miR-199a-5p also correlated with genes related to obesity and inflammation such as *PRDM16*, *ADAMTSL3, ZNF300* and *TAB3* (**Table S7**). Single-nucleotide polymorphisms near *ADAMTSL3* have previously been reported to be associated with lean body mass ^26^, while *ZNF300* plays a role in liver lipid metabolism ^27^ and *TAB3* is associated with pancreatic inflammation ^28^.

Another previous study also identified miR-199a-5p to be associated with Klotho (an anti-aging protein) and to activate downstream signalling pathways leading to fibrosis and inflammation in kidney cells ^29^. In our study, we identified the age-related gene *RBPMS* ^30^ to be associated with miR-199a-5p in human islets (**Table S7**). Some of the top-10 mRNAs, which showed significant negative correlation and contain conserved binding sites to miR-199a-5p are islet survival genes such as *JAG1* ^31^ and *TGFB2* ^32^ (**Figure 3A**).

MicroRNA-214-3p has previously been suggested to play a role in metabolic function and obesity. Serum miR-214 expression levels were lower in obese (especially in insulin-resistant) individuals compared to healthy individuals ^33^, whilst overexpression of miR-214 in hepatocytes ^34^ suppressed gluconeogenesis. Here, we identified miR-214-3p to be associated with *SORCS1* (**Figure 3B**), which is associated with insulin secretion ^35^, and metabolic genes such as *TPCN2* and *HFE* (**Table S8**). In human islets *TPCN2* is important for glucagon secretion and calcium elevation ^36^, while *HFE* expression is associated with iron content, insulin secretion and glucose desensitization in islets ^37^. Previous studies have also implicated miR-214 in the ageing of the liver ^38^ and in the control of mouse pancreas development ^39^. In this study, we observed that miR-214-3p was associated with age-related genes such as *ABCC6* ^40^ (**Table S8**).

We (**Figure 2C**) and others ^41^ found miR-214-3p associated with donor sex in human islets. The long non-coding RNA X-inactivation specific transcript (XIST, required for inducing X-chromosome inactivation in females) has been shown to interact with miR-214-3p ^42^. MiR-214-3p is also negatively correlated with the X-chromosome located gene *AR* (**Table S8**), which controls glucose/energy homeostasis and metabolic dysfunction development in various tissues (including skeletal muscle, adipose, liver and pancreas) differently between males and females ^43^.

MicroRNA-214 and miR-199a-2 are expressed from the same microRNA cluster on chromosome 1 and contain shared regulatory features. This may explain their common association with donor sex, age and BMI in our study. Previous studies on the miR-214/-199 cluster have identified association with regulatory networks such as DNA hypermethylation in embryonal carcinoma cells ^44^ and mouse development, through interactions with Twist-1 ^45^, cell migration and proliferation ^46^. Moreover, a study also identified the miR-199a/214 cluster as a negative regulator of brown and beige fat development and thermogenesis in adipocytes by directly targeting *PRDM16* (as part of a *PRDM16*-peroxisome PGC-1α transcriptional network) ^47^. *PRDM16* is a transcription factor and histone H3 methyltransferase regulating a thermogenic gene program important for brown and beige adipocyte maintenance ^48^, but it is also required for islet development ^49^. In our analysis we also observed that *FIGN* was the most significantly negatively correlative gene to both miR-199a-5p and miR-214-3p (**Figure 3**). Although the function of *FIGN* is unknown in islets, *FIGN* encodes for an ATP-dependent microtubule-severing protein, and depletion of *FIGN* had been shown to improve injured axon regeneration ^50^. In human islets, miR-199a-5p and miR-214-3p may potentially bind to *FIGN* to promote cell regeneration.

Our analyses also confirmed previous studies findings, as well as identified new target mRNAs associated with sex, age or BMI (**Table S3**). *SPESP1* expression was observed to be lower in females, consistent with a previous study ^7^. Other candidates associated with donor sex and higher expression in male islets include *CDK1*, *TOP2A* and *NOX5* (**Table S3**). *NOX5* is the nearest neighbour to *SPESP1*.

A previous study had observed upregulation of *Top2a* in pancreatic islets in a model of ventromedial hypothalamic lesions ^51^. *NOX5* is a key mediator of human islet insulin secretion, however, in an over-nutrition-induced environment, *NOX5* is involved in beta-cell damage ^52^. Higher abundance of these genes (*TOP2A* and *NOX5*) in male islets, could be linked with the predominance of diabetes prevalence in males, as well as males being more susceptible to islet cell damage compared with females ^1^. Similar sex-differences in ER stress induced genes have been recently reported ^53^.

Other new candidate genes, which had significantly higher expression in female islets compared to male islets, include *BARHL1*, *RPS4XP13, ART5, APOB, NLRP11* and *NLRP12*. The mRNA *BARHL1* was significantly increased in female compared with male islets. Expression of *BARHL1* is important for migration and survival of cerebellar granule neurons^54^. Pertinent to our observation, the counterpart of *Barhl1* in *C.elegans, ceh-30* has an anti-apoptotic function in survival for sexually dimorphic cephalic male sensory neurons, regulated by sex determination pathways ^55^. Some of the identified sex-associated mRNAs (such as *BARHL1* and *NOX5*) also contain putative genetic variants at their regulatory regions in islets (**Table S6**).

Our observations of multiple target genes associated with donor age are similar to a recent study, which examined the associations between the RNA-seq profiles of 37 laser-captured pancreatic islet microsections from different donors without diabetes with donor age (from 1 to 81 years old) ^15^. Our analyses identified *CDKN2A*, *PAX5, SLC24A2*, *RBFOX1*, *SIX3* and *CST2* to be upregulated with age, while genes *HIST1H3E*, *FGF23*, *SOX2*, *CETP* and *BUB1B* were downregulated with increasing age. The age-associated gene *CDKN2A* has previously been studied in islets ^56^; *CDKN2A* is an important cell cycle regulating gene in pancreatic beta-cells ^56^ and GWAS have identified it to be significantly linked to T2D ^57^. Our study also identified genes which have not been reported to be associated age, such as *ADTRP* and *PGM5P4-AS1*. *ADTRP* is involved in maintaining cell and tissue homeostasis in other tissue types (such as adipose tissue and liver) ^58^ and also contain putative genetic variants in islets (**Table S6**).

The analyses of islets genes associated with BMI (**Table S5**) identified genes involved in obesity, metabolism and inflammation such as *GUCY2C*, *ZNF536*, *MYBPC1*, *DOK2*, *BCL11B* and *AMY2A* that negatively correlated; and *GALNTL6, CASQ2* and *CLDN18,* which positively correlated with BMI. Our study also identified genes that have been reported to be associated with BMI (*GYG1P3* and *RASAL1*).

*GYG1P3* is a pseudogene of Glycogenin 1 (*GYG1*), which is expressed across the skeletal muscle, pancreas and brain, involved in glycogen storage in muscle and glycogen synthesis in muscle and heart ^59^. Chronic hyperglycaemia in a mouse model of human neonatal diabetes ^60^ results in marked glycogen accumulation in islets, and increased apoptosis in β-cells. *RASAL1* has been previously reported to inhibit cell growth of a hepatic tumor ^61^. Putative genetic variants around the regulatory region of *RASAL1* in human islets are present (**Table S6**).

This report presents analyses of microRNA data from a large number of human islet samples, which is a major strength of this study. The study also provides a diverse coverage of different age (range from 16 to 69 years old), and BMI (17.9 to 57.5 kg/m^2^), with similar distribution of female and male donors. We have incorporated and provided analyses after adjustments for the other traits, multiple testing, and machine-learning resampling approach (to eliminate sampling bias). Our study also identified multiple new targets associated with sex, age and BMI in human islets. In addition, a strength of this study is that we were able to compare microRNA and mRNA profiles in the same sub-set of human islet donors.

In our correlative analyses, we identified that the sex, age and BMI microRNAs were associated with mRNA genes involved in islet development, survival and function. We identified different metabolic (e.g. *PRMDM16, TPCN2, HFE, ADAMTSL3* and *TAB3)*, aging/senescence (e.g. *ABCC6* and *RBPMS*) or important pancreatic endocrine islet (e.g. *TGFB2, JAG1, SORCS1, TPCN2* and *HFE*) related genes associated with the sex, age and BMI microRNAs (miR-199a-5p and miR-214-3p).

In our functional studies, we observed that the inhibition of miR-199a-5p and miR-214-3p slowed down the shortening of RTL during cellular ageing represented by replicative cellular senescence (marker of senescence ^18^) in human islet-derived cells. Our observation aligns with previous studies which have similarly identified inhibition of miR-199a-5p reduced cellular senescence in vascular smooth muscle cells ^62^, and restored mesenchymal stem cells from senescence ^63^. MicroRNA-214-3p inhibition in past literature have also shown to promote bone formation in an ageing (ovariectomy, OVX) mice model ^64^, and could enhance proliferation and migration in endothelial cells ^65^. Our functional studies imply that targeting miR-199a-5p and miR-214-3p could retard age-related alterations in human islets.

Overall, our study reports microRNAs (and their targets) associated with donor traits, and also presents sub-analyses for potential microRNA-mRNA associations in human islets. Our analyses of these data provide a cautionary note to studies of specific microRNAs (and mRNA species) that may be driven by donor trait(s).

A limitation of this study is that our islet donors represent samples from human islet isolation sites within Australia. Comparison of our findings with human islets obtained from other islet isolation centres would help us understand ethnic and/or processing ^66^ differences. Our studies are limited to associations since methods of replicating ageing or obesity *in vitro* are difficult. Future studies at a single-cell scale will help identify specific pancreatic cell types, which are different in transcriptomic profile between certain donor traits. Single-cell RNA-seq analyses of pancreatic islets have been used to identify differences for individually characterized pancreatic cell types associated with donor age ^16,67^. We previously assessed microRNAs and candidate target (mRNA) gene transcripts at single cell resolution ^68,69^. Current technological limitations in single-cell omics for profiling both microRNA and mRNAs in human islets prevent the much-wanted datasets for single-cell miRNA and mRNA profiling. Although single-cell sequencing provides the resolution to assess cellular heterogeneity, such studies often lack the desired robustness across many donors. Future technological developments in analysing microRNAs and mRNAs from same (half-)cells may enhance our observations of the association of islet microRNAs with donor traits.

## 4. Materials and Methods

### 4.1. Human islets

Isolated cadaveric human islets (**Table S1**) from donors without diabetes were obtained through the Australian Islet Transplantation Program (at Westmead Hospital, WIMR, Sydney, and SVI, Melbourne), following the human research ethics committee (HREC) approvals from Sydney Local Health District.

### 4.2. RNA isolation, quantification, and quality

Total RNA isolation and quantification for concentration and RNA integrity number (RIN) performed are described previously ^70–72^.

### 4.3. Nanofluidics-based TaqMan® real-time PCR on 754 validated human microRNAs

Nanofluidic-based TaqMan® OpenArray™ human microRNA panel was used to quantify 754 validated human microRNAs on the QuantStudio™ 12K Flex platform (Thermo Fisher Scientific, Waltham, MA) ^73^. Details of PCR analysis are described in our previous work ^71^. MicroRNA transcript abundance was log2 transformed for statistical analyses.

### 4.4. Bulk RNA-sequencing and analysis

RNA-sequencing on human islet samples (GSE152111) was performed on the HiSeq4000 platform at 150 paired-end reads ^72^. RNA-sequencing analysis was carried out using the Strand Next-generation sequencing (NGS) version 2.5 software ^72^. Samples were aligned to human hg38 transcriptome and genome together (novel splice variants) with Ensembl annotation. Quality trimming and filter (base quality < 20 removed) was applied after. An average of 29 million clean reads per sample was generated. Aligned reads of all samples were quantified and normalized with threshold normalized count set at one, using DEseq. Two of the 66 islet samples clustered with the opposite sex group (on unsupervised Euclidean average cluster analysis performed on the X and Y chromosome genes across all samples), therefore were removed for further analysis. A donor sample (H149) was duplicated in RNA-sequencing. The transcriptome profile of the duplicates was highly correlative (Pearson r=0.99) to each other. Average of the duplicates from same donor sample (H149) was calculated and used for analysis.

### 4.5. Cell culture

Human primary islet derived cells were maintained in serum-containing medium (CMRL, with 1% GlutaMAX™, 10% fetal bovine serum (Thermo Fisher Scientific, Waltham, MA), human Epidermal Growth Factor (hEGF) 10 ng/ml (Sigma-Aldrich, St Louis, MO), 100U/mL penicillin and 100μg/mL streptomycin (Thermo Fisher Scientific, Waltham, MA). Cells were maintained in an incubator at 37°C and with humidified 5% CO2 in air. For microRNA knock-down experiments, islet-derived cells were cultured in six-well tissue culture plates and 50nM of each of the two LNA power inhibitors targeting the two microRNAs ((miR-199a-5p andmiR-214-3p) were added to one well of the plate. Untreated cells were used as controls. Once cells were confluent in their respective wells, we trypsinised them, and harvested 50% of the cells for RNA and DNA, while remaining 50% were added onto a new well. To induce replicative cellular senescence, the islet derived cells went through multiple passages for up to 31 days, with the addition of LNA power inhibitor for the entire duration. MicroRNA inhibition (knockdown) was carried out using microRNA LNA Power LNA inhibitors (Qiagen, Hilden, Germany) as previously described in detail in our protocol paper ^74^

### 4.6. Individual microRNA RT-qPCR

Individual microRNA (U6 (internal control), miR-199a-5p and miR-214-3p) expression was quantified using TaqMan® qPCR chemistry on the ViiA7 platform (Thermo Fisher Scientific, Waltham, MA), as previously described in detail in our publication ^73^. Quantitative expression levels (Cycle threshold (Ct) values) of miR-199a-5p and miR-214-3p were normalized to internal control U6. Normalized Ct values was used to calculate the transcript abundance using formula “=2^(39-Ct^ ^value)^”, where “39 Ct value” is the limit of detection ^75^.

### 4.7. Relative telomere length quantification

DNA was isolated using Qiagen kit (Qiagen®, Hilden, Germany) and used to quantify relative telomere length as we have previously described in detail ^18^. Relative telomere length was quantified using primers for telomere ((A): CGGTTTGTTTGGGTTTGGGTTTGGGTTTGGGTTTGGGTT, Telomere (B): GGCTTGCCTTACCCTTACCCTTACCCTTACCCTTACCCT) and separately primers for human β-globin gene (hbg1: GCTTCTGACACAACTGTGTTCACTAGC, hbg2: CACCAACTTCATCCACGTTCACC) used as single-copy gene. Telomere and single-copy gene (human β-globin) were separately quantified via SYBR green qPCR for each sample in triplicates. Here, relative telomere length (RTL) is presented in ΔCt and calculated for each sample using the “(average Ct of hbg-average Ct of telomere)”. Percent change compared to the baseline (Day 0) RTL was calculated and plotted using GraphPad Prism.

### 4.8. Statistical analysis

Statistical analyses were performed using R software (ver. 3.6.2-3), SPSS Statistics 27, Microsoft Excel (ver. 2016) or GraphPad Prism 8.4.1. L1-Penalized regression (Lasso) (PLR) analysis was carried out using logistic (for sex) or linear (age or BMI) regression workflow, followed by bootstrapping (n=1000 iterations) using R packages penalized (0.9.51) and glmnet (4.0-2) as described previously ^70,71^. Kolmogorov-Smirnov test was used to check for normal distribution using SPSS. F-test was performed to check for variance between male and female. Two-tailed Mann–Whitney U test was used to calculate p-values, computed with no ties in R using wilcox.test() for non-normally distributed data. Two-tailed Student’s (for equal variance) and Welch’s (for unequal variance) t-test was used for normally distributed data. Benjamini-Hochberg approach was used for False Discovery Rate (FDR) multiple testing. Statistical adjustment for sex, age and/or BMI were calculated with multiple (linear) regression model using the lm() function in R. Kruskal-Wallis test for trend analysis was performed on GraphPad Prism. Significance classified as p-value ≤0.05.

### 4.9. In silico analysis

The predicted potential binding sites located at and near the gene targets to the microRNAs of interest were obtained using computational web tool TargetScan Human ver. 8 ^76^.

### 4.10. Human islet Expression Quantitative Trait Loci (eQTLs) database analysis

The Translational Human Pancreatic Islet Genotype Tissue-Expression Resource (TIGER) database (http://tiger.bsc.es/) was used to search for significant expression quantitative trait loci (eQTLs) in the genes most significantly and negatively associated with sex, age and BMI microRNAs.

### 4.11. Human T2D Genome-Wide Association Study (GWAS) database analysis

The T2D Knowledge Portal (https://t2d.hugeamp.org/) was used to search for genetic variants in the genes most significantly associated with donor sex, age and BMI in islets identified in this study.

### 4.12. Enrichment pathway analysis

Pathway analyses on predicted target mRNAs of the sex, age and/or BMI associated microRNAs (TargetScan Human ver.8) were filtered using a reference list of beta-cell expressed genes (from E-GEOD-20966). Over-representation analysis was performed using Panther, Pantherdb.org ^77^, and with Gene Ontology: Biological processes (GOBP) classifications using beta-cell expressed mRNAs as background set. Fishers’ exact test and FDR correction for multiple testing was used.

## Acknowledgements

The research presented herein has been funded through grants from the Australian Research Council Future Fellowship (FT110100254), the Juvenile Diabetes Research Foundation (JDRF) Australia T1D Clinical Research Network (JDRF/4-CDA2016-228-MB) to A.A.H. This study was supported by the Marian and E.H. Flack Trust (to W.J.H; FLACK). A.A.H is also supported through a Visiting Professorship from the Danish Diabetes Academy, funded by the Novo Nordisk Foundation, grant number NNF17SA0031406 (2016-18 and 2019-23). W.K.M.W. acknowledges previous support from the Australian Postgraduate Award, University of Sydney, JDRF Australia PhD top-up award, and current funding through JDRF Australia/Helmsley Charitable Trust. M.V.J. was supported through a JDRF USA advanced post-doctoral award (3-APF-2016-178-A-N) and a transition award from JDRF, USA. The St Vincent’s Institute receives support from the Operational Infrastructure Support Scheme of the Government of Victoria. R.C.W.M. acknowledges support from the RGC Theme-based Research Scheme (T12-402/13N) and Research Impact Fund (R4012-18), the Focused Innovation Scheme, and Faculty Postdoctoral Scheme of the Chinese University of Hong Kong. Postdoctoral Fellowship Scheme, The Chinese University of Hong Kong (to G.J). A.E.S. was supported through a Danish Diabetes Academy post-doctoral grant, supported by the Novo Nordisk Foundation. Infrastructure support from the Faculty of Medicine & Health, University of Sydney; Australia, School of Medicine, Western Sydney University, Australia; Western Sydney University, Ingham Institute, Liverpool; Australia and the Rebecca L. Cooper Medical Research Foundation, is acknowledged. The support of all surgical teams and members contributing to the acquisition of research tissue samples, all organ donors, and supporting family members is gratefully acknowledged.

## Author contributions

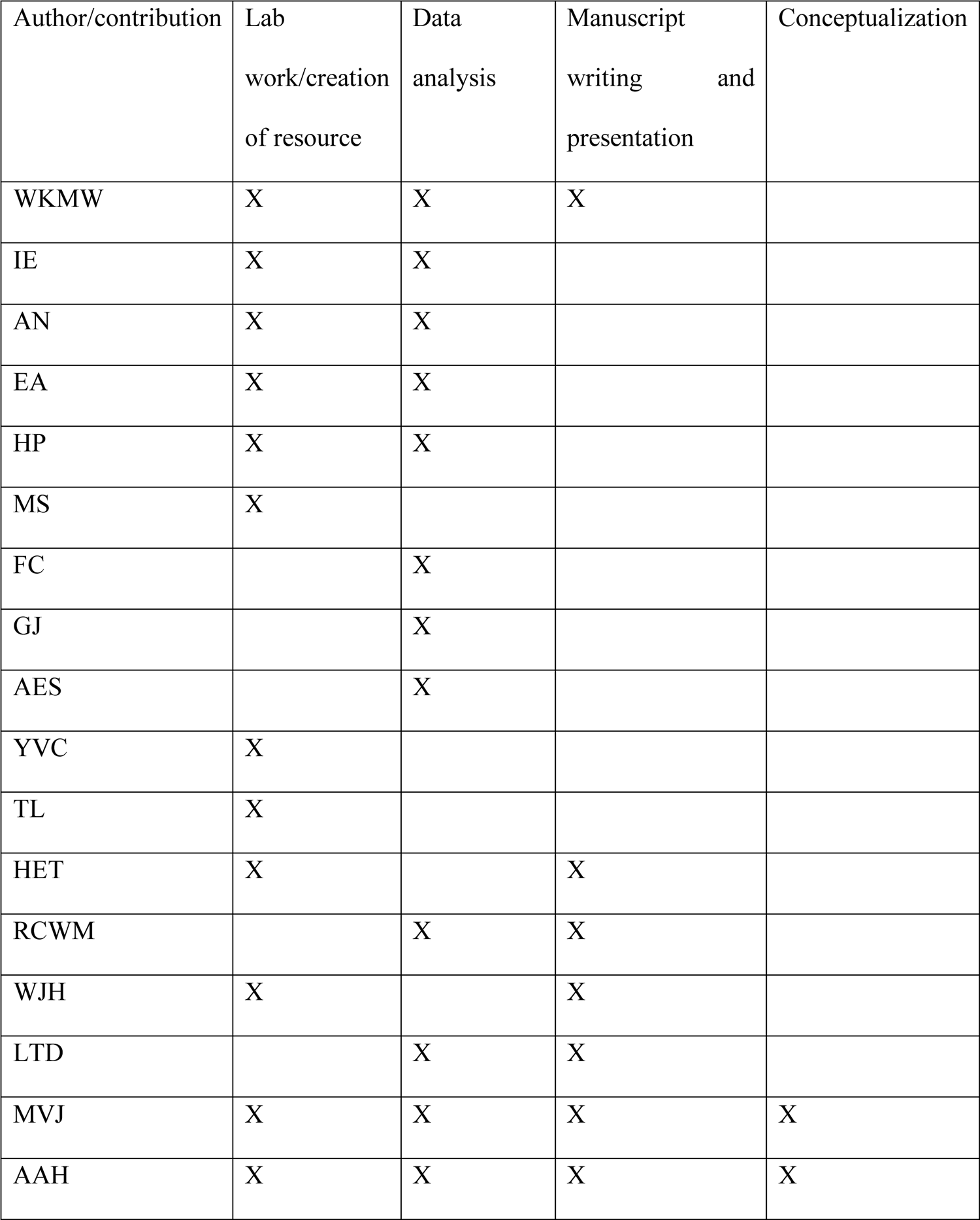

## Data availability

Human islet RNA-sequencing data used are available on the Gene Expression Omnibus (GEO) database, accession number GSE152111.

## Tables

Excel Worksheets attached

---

**Figure S1.**
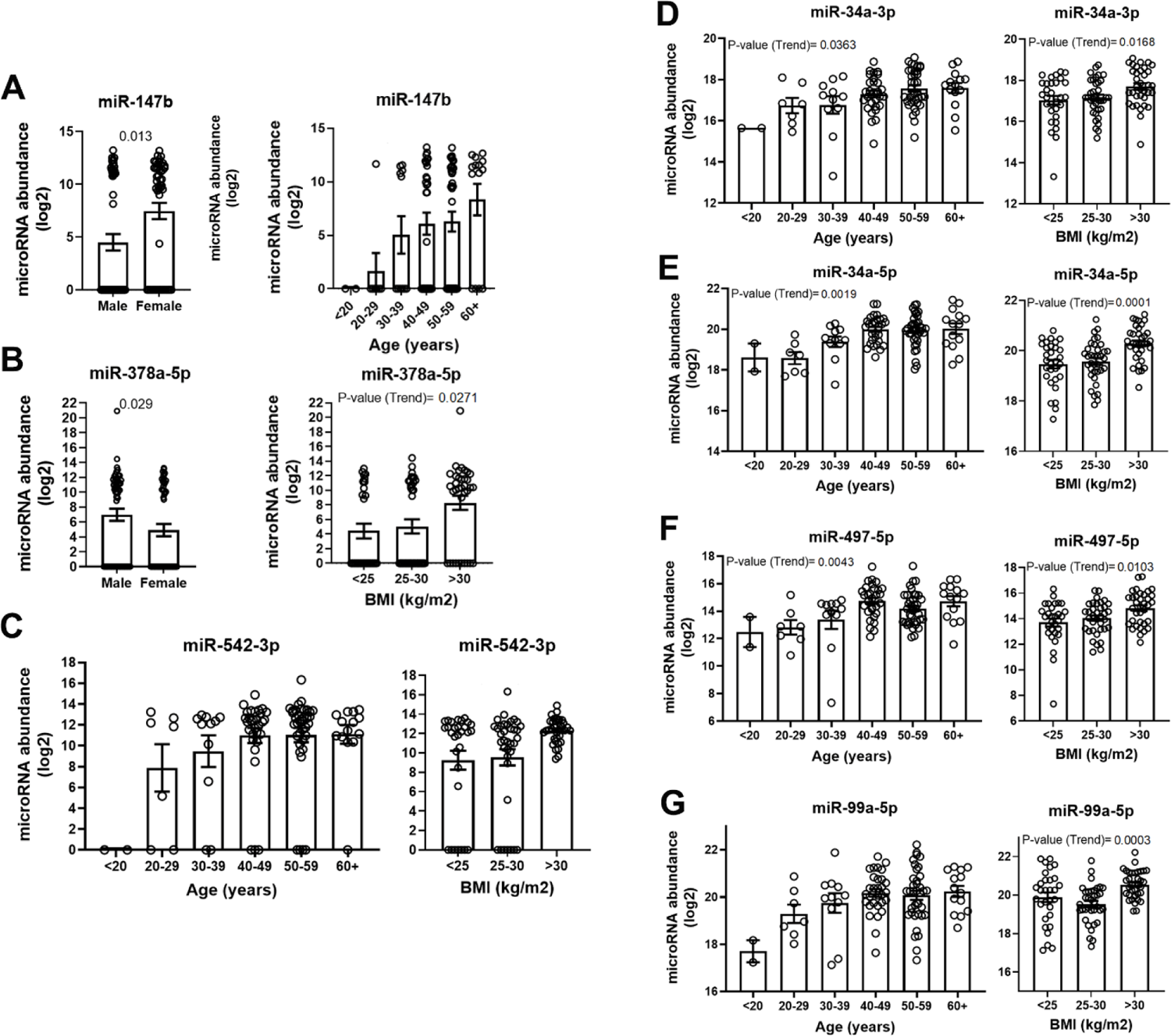
Human islet microRNAs associated with sex, age and BMI. MicroRNA transcript abundance (log2) for **(A)** miR-147b on sex and age; (**B**) miR-378a-5p/miR-378 on sex and BMI; (**C**) miR-542-3p, (**D**) miR-34a-3p, (**E**) miR-34a-5p, (**F**) miR-497-5p and (**G**) miR-99a-5p on age and BMI. MicroRNA quantification was performed using TaqMan rt-qPCR. The adjusted (for age and BMI) p-value between male and female (sex) comparison are presented. Trend analysis (using Kruskal-Wallis test) was performed to calculate p-value (trend) presented (in figures) for age or BMI. Significant trend values are provided. Mean ± SEM are presented.

**Figure S2.**
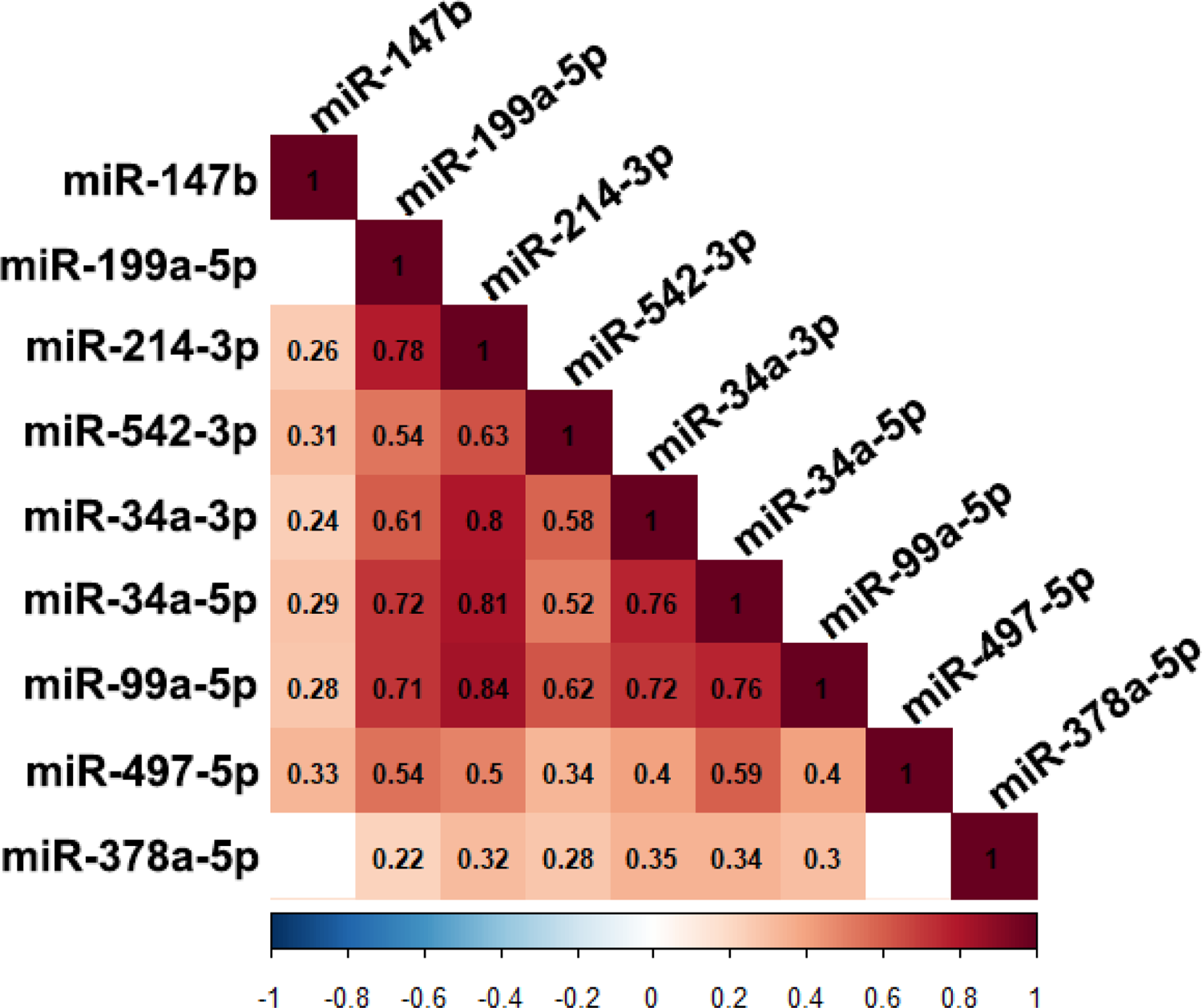
Human islet sex, age and BMIC associated microRNAs associated correlation. Correlation between sex, age and BMI associated microRNAs (miR-147b, miR-199a-5p, miR-214-3p, miR-542-3p, miR-34a-3p, miR-34a-5p, miR-99a-5p, miR-497-5p and miR-378a-5p) in human islets (n=101). The spearman correlation rho coefficients are presented in black text in each square. Only the significant correlations (Spearman p ≤ 0.05) are presented with a coloured (as depicted in the scale bar) in the grid displaying their Spearman correlation (rho, r) coefficient.

**Figure S3.**
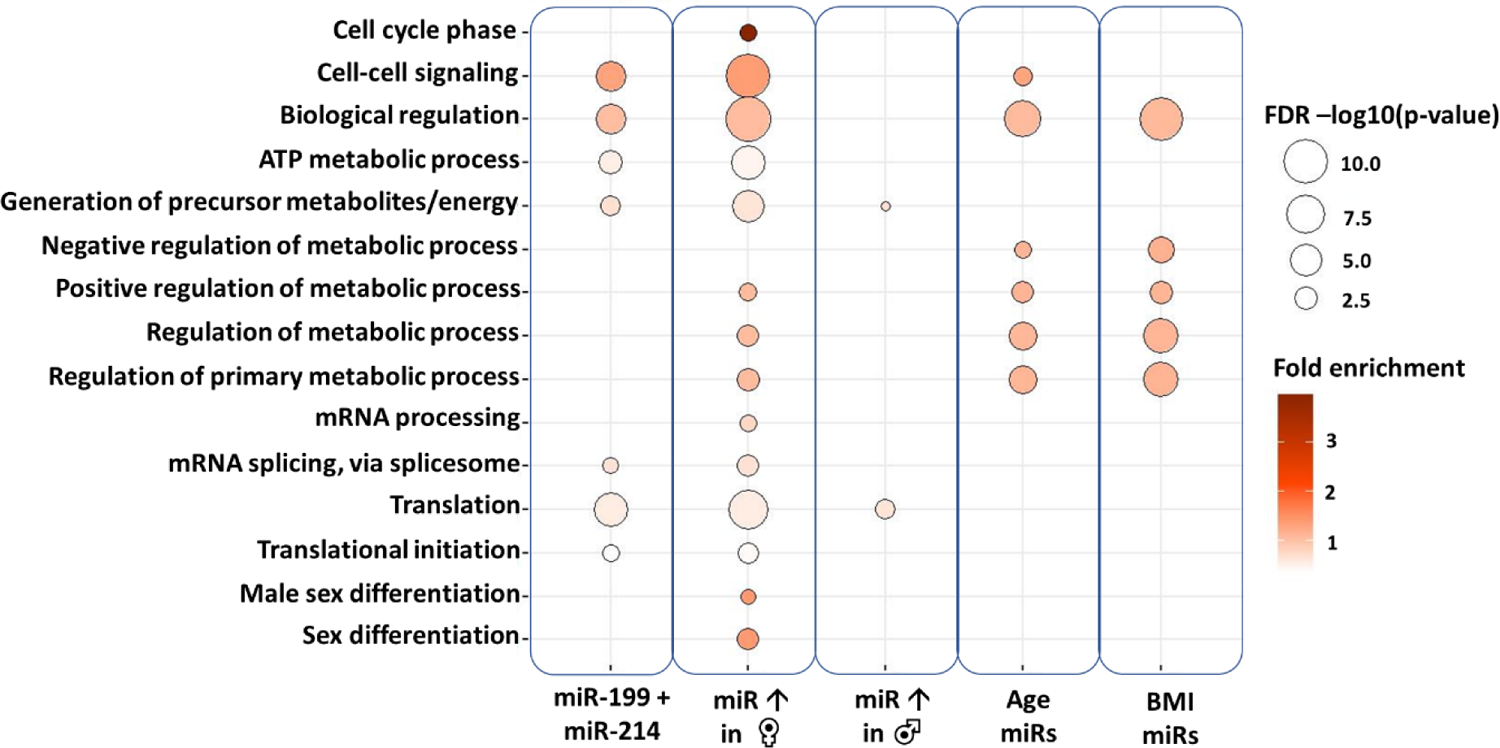
Sex, age and BMI associated islet microRNAs are involved with sex differentiation, transcriptional, translational, cell cycling and metabolic enriched pathways. GOBP Enrichment pathway analyses performed on the significant islet microRNAs associated with donor sex, age and BMI. Significant selected cell cycle and signalling, metabolic, mRNA, translational and sex differentiation pathways to the sex associated and/or BMI islet microRNAs are presented. Age miRs: represents the top 11 most significant microRNAs (after adjustment; miR-34a-5p, −34a-3p, −29b-3p, −29c-5p, −206, −147b, −379-5p, −339-3p, −642a-5p, −16-5p, −376c-3p) associated with donor age. BMI miRs: represents the top 11 most significantly microRNAs (after adjustment; miR-155, −34a-5p, −130a-3p, −18a-5p, −190a-5p, 193a-5p, 7f-2-3p, 19a-3p, −497-5p, 148b-5p and −27b-3p) associated with donor BMI. MiR ↑ in ♀: represents the microRNAs (miR-147b, −130b-3p, −218-5p, −199a-5p, −544a, −214-3p, −7c-5p and −138-5p; after adjustment) that were significantly higher in female compared to male islets. MiR ↑ in ♂: represents the microRNAs (miR-205-5p and −378a-5p; after adjustment) that were significantly higher in male compared to female islets. The size of the bubble displays the False Discovery Rate (FDR) log10 transformed p-value. The shades of red displays the fold enrichment for each pathway to the specific set of microRNAs.

**Figure S4.**
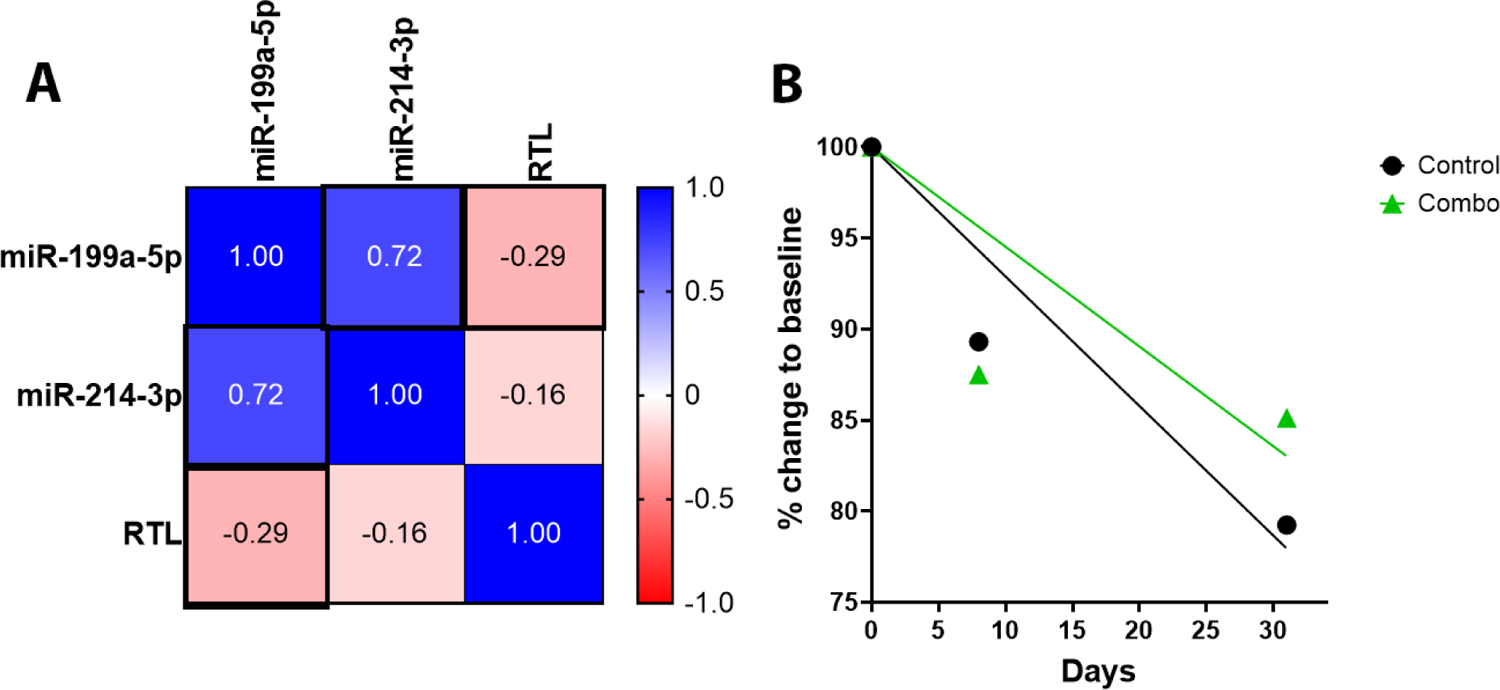
Human islet miR-199a-5p and −214-3p association with relative telomere length. (**A**) Spearman correlation matrix of the expression levels of miR-199a-5p, miR-214-3p and relative telomere length (RTL) measured in different passage time points (up to 16 time points, in n=8 different islet donor samples). Statistically significant (P≤0.05) correlation highlighted with bold borders. The Spearman rho coefficients are presented within each box and by color (from blue to red, displaying positive to negative correlation). (**B**) The percentage of relative telomere length change to baseline of islet-derived preparations (n=3 different islet donor samples) (Y-axis) at different cell culture passage time points (day 8 and day 31, X-axis). Average of the different islet-derived cell preparation are presented for control (black dot) and combination of miR-199a-5p and −214-3p inhibition (combo, green triangle). Slope line of best fit are presented in black line (for control) and green line (for combo).

